# Effects of intensified training with insufficient recovery on joint level and single muscle fibre mechanical function: The role of myofibrillar Ca^2+^ sensitivity

**DOI:** 10.1101/2023.10.13.562290

**Authors:** Olivia P. Roussel, Christopher Pignanelli, Emma F. Hubbard, Alexandra M. Coates, Arthur J. Cheng, Jamie F. Burr, Geoffrey A. Power

## Abstract

Intense exercise training with insufficient recovery is associated with reductions in neuromuscular performance. However, it is unclear how single muscle fibre mechanical function and myofibrillar Ca^2+^ sensitivity contribute to these impairments. We investigated the effects of overload training on joint-level neuromuscular performance and cellular-level mechanical function. Fourteen athletes (4 female, 10 male) underwent a 3-week intensified training protocol consisting of ∼140% of their regular training hours with three additional high-intensity training sessions per week. Neuromuscular performance of the knee extensors was assessed via maximum voluntary contraction (MVC) force, electrically evoked twitch contractions, and a force-frequency relationship. Muscle biopsies were taken from the *vastus lateralis* to assess single fibre mechanical function. Neither MVC force nor twitch parameters were altered following intensified training (all *p>*0.05), but a rightward shift in the force-frequency curve was observed with a 6-27% reduction in force at low-frequencies (5-20Hz, all *p<*0.05). In single fibres, maximal force output was not reduced following intensified training, but there was a rightward shift in the force-pCa curve driven by a 6% reduction in Ca^2+^ sensitivity as indicated by a lower pCa_50_ value (i.e., higher [Ca^2+^]) across fibre types (Pre=6.477±0.157, Post=6.088±0.480, *p<0.05*). These data indicate intensified training leads to impaired Ca^2+^ sensitivity at the single fibre level, which in part explains impaired neuromuscular function at the joint level during lower frequencies of activation. This is an important consideration for athletes, as performance is often assessed at maximal levels of activation, and these underlying impairments in force generation may be less obvious.

**New & Noteworthy:** Intense exercise training with insufficient recovery leads to impaired muscle contractile performance. These impairments often manifest at lower frequencies of muscle stimulation, termed prolonged low-frequency force depression. Impaired myofibrillar calcium sensitivity has been suggested as a potential mechanism of prolonged low-frequency force depression. Our work shows that impaired calcium sensitivity of single muscle fibres coincided with joint level prolonged low-frequency force depression after intense exercise training with insufficient recovery.

## Introduction

Exercise training is well-established to promote health benefits and improve sport performance (15). Training adaptations are the result of acutely stressing physiological systems by manipulating the duration, frequency, and/or intensity of exercise (14, 15). However, when an increase in training load (i.e., intensified training) is not followed by adequate recovery, the benefits of exercise for both sport performance and overall health may be reduced (22, 26, 32, 38). While the effects of intensified training with insufficient recovery on the neuromuscular system are thought to play a role in the acute impairment of sport-specific tasks (6, 12), the underlying mechanisms are poorly characterized (8).

Strength can be assessed by performing an isometric maximum voluntary contraction (MVC), which is commonly used to measure changes in neuromuscular performance after a period of intensified training (6, 32). However, MVC force has previously been shown to not decrease following intensified training with insufficient recovery in both team sport athletes (6) and military conscripts (32). These findings indicate that MVC, or other assessments requiring short bursts of maximal effort, may not be the most appropriate to assess the effects of intensified training with insufficient recovery on neuromuscular performance. This may be in part because short, maximal efforts are associated with higher motor unit discharge frequencies (∼20-50Hz (36)), which could conceal impairments in neuromuscular function at lower motor unit discharge frequencies. It is well-established that muscle force production can be depressed hours to days after exercise at low- relative to high-frequency electrical stimulation even with no changes in MVC force (13, 37). The prolonged reduction in force at low frequencies of stimulation, relative to high-frequency stimulation has been termed prolonged low-frequency force depression (PLFFD;(1)), and may provide further insight into how intensified training with insufficient recovery influences neuromuscular performance. Mechanistically, PLFFD may involve a decrease in myofibrillar Ca^2+^ sensitivity and/or reduced Ca^2+^ release from the sarcoplasmic reticulum (9).

Myofibrillar Ca^2+^ sensitivity determines the amount of force that can be produced by the contractile proteins for a given myoplasmic free [Ca^2+^] ([Ca^2+^i]), and reductions in sensitivity may impair force production at submaximal levels of [Ca^2+^i] (23). Both a decrease and increase in myofibrillar Ca^2+^ sensitivity has been reported following bouts of prolonged cycling or repeated high-intensity cross-country skiing in world-class athletes (17, 21), possibly as a result of redox modifications of specific residues on proteins located in the contractile apparatus (28). While impaired Ca^2+^ release from the sarcoplasmic reticulum is implicated as a mechanism of PLFFD in skeletal muscle (4, 7, 31, 41), previous work in rodent muscle has also shown PLFFD can be driven by either impaired Ca^2+^ release or reduced myofibrillar Ca^2+^ sensitivity (4, 7, 40). The relative weighting of impaired Ca^2+^ release and myofibrillar Ca^2+^ sensitivity contributing to PLFFD can be altered when an antioxidant enzyme, superoxide dismutase 2 (SOD2) is overexpressed (4). Given endurance training increases muscle antioxidant defence systems including SOD2 expression (e.g., (33) and reviewed in (35)), it is possible the impaired neuromuscular performance after intensified training with insufficient recovery in endurance-trained athletes may occur because of reductions in myofibrillar Ca^2+^ sensitivity.

The purpose of the present study was to characterize how intense exercise training with insufficient recovery affects neuromuscular function using joint-level neuromuscular assessments and single muscle fibre mechanical testing to assess myofibrillar Ca^2+^ sensitivity. It was hypothesized that: 1) intensified training would result in PLFFD, with minimal effects on both maximal voluntary force and high frequency electrically stimulated force and 2) there would be a reduction in myofibrillar Ca^2+^ sensitivity following intensified training.

## Methods

### Study overview

Fourteen tier 2 to tier 3 endurance athletes ((25); 4 female, 10 male) completed this study. All participants were training a minimum of 7h a week prior to study commencement (Table 1). The methods for the outcomes in Table 1 as well as the impact of intense training and recovery on sport performance in participants classified as overreached can be found in a related work (11). Most training was endurance-type (e.g., running, swimming, cycling, rowing), and some participants continued supplementary resistance-type exercise throughout the study. This study was part of a larger study examining the cardiovascular, metabolic, and neuromuscular adaptations to intensified training. The study was reviewed and approved by the University of Guelph research ethics board (REB#21-11-017) and all participants provided written informed consent. An overview of the study design and relevant measures in this study can be found in Figure 1.

**Figure 1.**
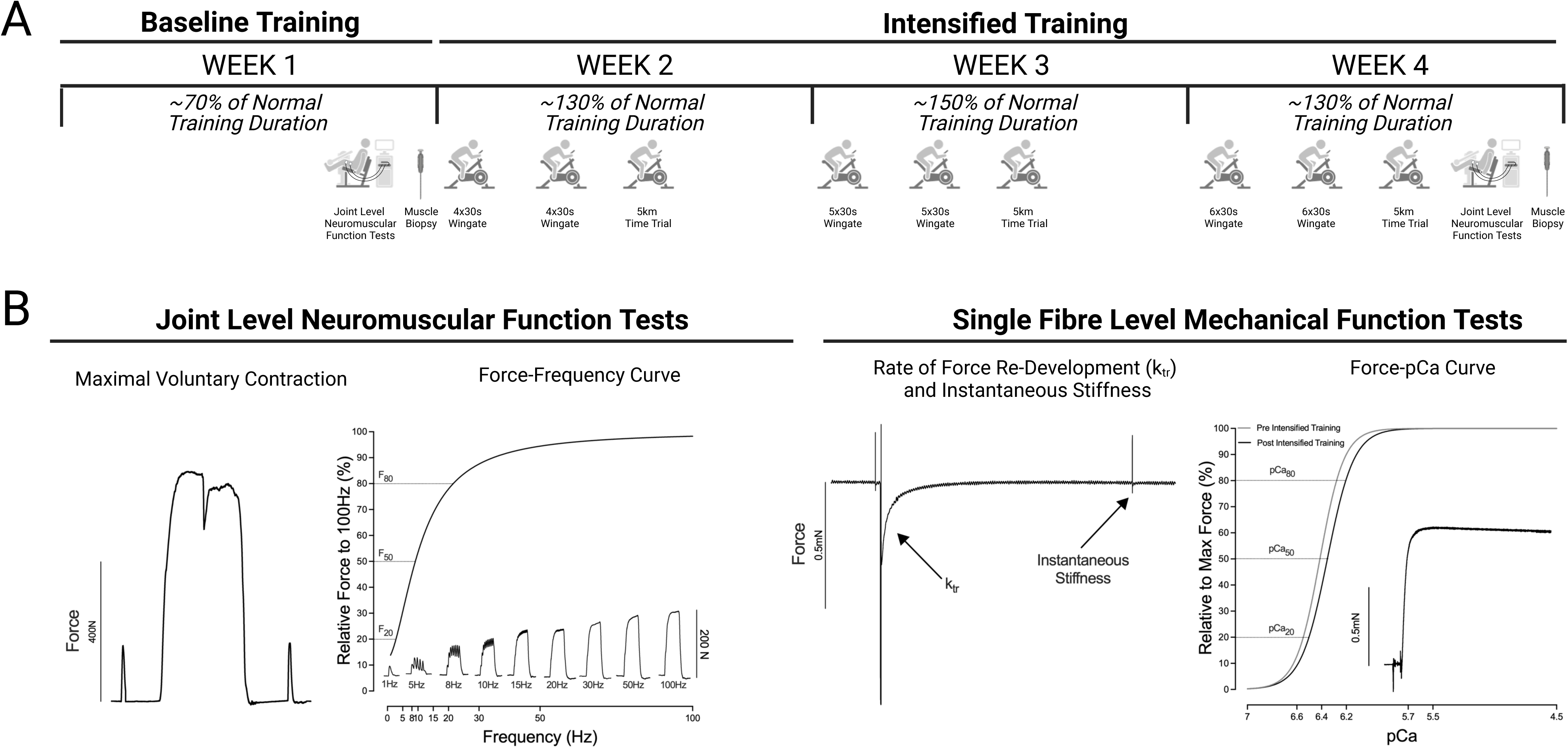
(A) Overview of the training study. Joint level neuromuscular tests and muscle biopsies were obtained at the end of the first week of training with ∼70% of the normal training duration. Participants then underwent three weeks of intense exercise training by increasing the exercise duration to ∼130%, ∼150%, and ∼130% their normal training hours, which included three high-intensity sessions composed of two Wingate sessions and a 5km time trial. Joint level neuromuscular tests and muscle biopsies were obtained at the end of the third week of intensified training (i.e., WEEK 4). (B) Representative tracings of the joint level neuromuscular function tests: maximal voluntary contraction with the interpolated twitch technique and a curve fitting procedure using non-linear regressions analysis to determine the frequency to elicit 20, 50, and 80% of the force produced at 100Hz during the force-frequency test (inset). The dotted lines represent the frequency needed to evoke the 20% (F_20_), 50% (F_50_), and 80% (F_80_) of the force produced at 100Hz. Single fibre level mechanical function tests: force production, rate of force re-development (k_tr_) and instantaneous stiffness tests at a pCa concentration of 4.5 and a representative fitted trace of the force-pCa curve of a single muscle fibre showing a shift to the right (i.e., decreased Ca2+ sensitivity) pre and post training. The dotted lines represent the pCa concentration needed to evoke the 20% (pCa_20_), 50% (pCa_50_), and 80% (pCa_80_) of the peak force. The figure was in part created with BioRender.com.

**Table 1.** Participant Characteristics.

|  | Male (n=10) | Female (n=4) | Total (n=14) |
| --- | --- | --- | --- |
| **Age (years)** | 28±5 | 25±6 | 27±6 |
| **Height (cm)** | 181±6 | 165±9 | 176±10 |
| **Body mass (kg)** | 82±9 | 59±7 | 75±14 |
| **Baseline training (hours)** | 8.3±3.0 | 10.9±3.1 | 9.0±3.1 |
| **Aerobic Peak Power** |  |  |  |
| Absolute (Watts) | 406±63 | 283±43 | 371±81 |
| Relative (Watts/kg) | 4.96±0.67 | 4.84±0.70 | 4.93±0.65 |
| **$\dot{V}O_2$ peak** |  |  |  |
| Absolute (L/min) | 4.61±0.69 | 2.84±0.75 | 4.10±1.07 |
| Relative (mL/min/kg) | 56.2±6.6 | 48.6±11.8 | 54±8.7 |
| **$\dot{V}O_2$ at first ventilatory threshold** |  |  |  |
| Absolute (L/min) | 3.22±0.65 | 1.95±0.33 | 2.85±0.82 |
| Relative (mL/min/kg) | 39.2±6.9 | 33.3±4.6 | 37.5±6.7 |
| Percent of $\dot{V}O_2$ peak (%) | 69.5±6.6 | 70.7±14.2 | 69.8±8.8 |

Participants completed a total of four weeks of training with this protocol (Fig. 1A; (11)). In the first week, participants were asked to perform ∼50% of their normal training hours prior to the study and were familiarized with the joint level neuromuscular tests. Participants then underwent 3 weeks of intensified training with the aim of accumulating ∼150% of their normal training hours. During the intensified training period, participants were instructed to perform their regular training sessions, and partake in 3 additional supervised high-intensity training sessions each week. These sessions consisted of two Wingate sessions (30s sprints at a load of 7.5% body mass with 4min of active recovery between intervals) that increased in repetitions from 4×30s Week 2, 5×30s Week 3, to 6×30s Week 4, and a 5-km time trial on a cycle ergometer (Velotron, Racermate, USA). Only the high-intensity sessions were performed in the laboratory, and the remainder of participant training sessions were recorded via an online application (Strava, Inc). In total, the average training duration for each week is as follows: Week 1 training duration was ∼70% (6.4±0.6h) and Week 2, 3, 4 were ∼130% (11.8±0.8h), ∼150% (13.7±1.0h), ∼130% (12.0±1.2h) of the normal training hours prior to starting the study (Fig. 1A).

Joint level neuromuscular testing of the knee extensors and a skeletal muscle biopsy from the *vastus lateralis* were obtained only before (i.e., at the end of Week 1) and during the intensified training period (i.e., end of Week 4). Joint level neuromuscular testing was performed ∼24 hours after the last intense training session. Muscle biopsies of the vastus lateralis were obtained before the intensified training period and approximately 72h after the last 5-km time trial and/or intense training session. Participants were instructed to perform no more than 1h of low intensity exercise the day prior to each biopsy, and the exercise was matched for pre-and and post-testing. Menstrual cycle was not controlled in this study, however, measurements obtained herein were ∼4 weeks apart, thus, female participants were likely in similar hormonal phases for each testing period.

### Joint level neuromuscular testing

Knee extensor neuromuscular function was measured using electrically evoked twitch contractions, isometric MVCs with the interpolated twitch technique to assess voluntary activation, and a force-frequency curve (Fig. 1B). In the fasted state, participants performed resting metabolic and cardiovascular measurements (∼3h). Afterwards, participants consumed the same light meal 30 to 60min prior to the neuromuscular testing. Participants were allowed food and caffeine prior the neuromuscular testing to ensure they could perform maximally. Previous work supports a negligible effect of caffeine consumption for our primary outcome of PLFFD (29). Two electrodes (9cm wide x 15-20cm long), customised to leg girth, were coated in transmission gel and placed on the upper and lower right thigh and secured with tensor wraps and tape to ensure good contact. Participants were then fitted on a custom-built isometric dynamometer for subsequent testing. The right ankle of the participant was attached perpendicular to a calibrated load-cell (Model SML- 300; Durham Instruments, CAN) by a non-compliant cuff, with their knee joint at approximately 90°. Data was acquired through PowerLab 16/35 hardware and recorded on LabChart 8 software (ADInstruments, AU) sampled at 1 kHz. The current that elicited the maximal twitch force was found for each participant with a high-voltage stimulator (pulse width of 200µs) by stimulating participants until the twitch force no longer increased with increasing current (DS7AH; Digitimer, UK). This stimulation current was increased by 20% and used during the interpolated twitch technique to estimate voluntary activation, described below.

### Maximal voluntary contraction force and activation

The interpolated twitch technique was used to estimate voluntary activation (27). In brief, a twitch was delivered before, during, and after each MVC attempt. Voluntary activation was quantified by identifying the change in force when stimulation was delivered during the MVC (superimposed twitch force) and dividing it by the twitch force following the MVC (the potentiated twitch). The equation used to calculate voluntary activation was:

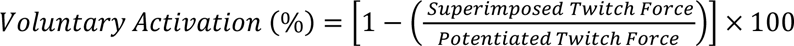

During each MVC, participants received strong verbal encouragement and visual feedback of force production on a computer monitor positioned approximately 1m from the participant. Guidelines were displayed to provide motivation to attain a higher maximum force with each attempt. The participants were given a minimum of 3 attempts separated by 3-5min of rest. Objective criteria to deem an MVC as maximal effort was 1) no further increase in force between attempts and 2) voluntary activation of ≥90%. Participants were given two additional attempts if they felt they could attain a higher force value after the initial 3 attempts. The maximal MVC force is reported as the highest 500ms average force from a single attempt.

### Force-frequency curve with electrically stimulated contractions

After the MVCs were completed, the current that elicited 30% of MVC force at 100 Hz stimulation was determined. Following 3-5mins of rest, 1s of stimulation (pulse width of 200µs) was delivered at the following frequencies, each separated by 20s: 1, 5, 8, 10, 15, 20, 30, 50, and 100Hz in that order to minimize the potentiating effects of higher Hz stimulation on low Hz force. Peak force, determined as the highest force reached during the contraction, was recorded at each frequency. One participant expressed discomfort during the stimulation at 100Hz to evoke 30% MVC at baseline testing and, as such, the relative intensity was reduced to 20% MVC for both timepoints for this participant.

### Muscle biopsies

Ten participants consented to providing resting muscle biopsies from the *vastus lateralis*. Baseline samples were obtained on the left leg and after the 3 weeks of intensified training, another biopsy was obtained on the right leg. Biopsies were performed under local anesthesia (2% lidocaine with epinephrine) using the Bergström technique. Samples were freed from blood and/or connective tissue, and a section of each sample was quickly placed into chilled skinning solution. See *Supplemental Information* (10.6084/m9.figshare.24231727) for skinning solution composition and all other solutions used. Subjects were asked to refrain from non-essential supplements (e.g., caffeine), alcohol, or other recreational drugs for at least ∼24h, exercise for ∼24h and food/liquids for 12h before each sampling.

### Single fibre testing procedure

Fresh muscle bundles were placed in chilled skinning solution for 30 minutes and maintained on ice. Gentle agitation was performed to ensure sufficient permeabilization in the skinning solution. The bundles were washed with fresh, chilled dissecting solution and gently agitated to remove any remaining skinning solution from the bundles. The bundles were then placed in a separate tube with the storage solution and stored for 24h at 4°C. See *Supplemental Information* (10.6084/m9.figshare.24231727) for all solution recipes. The tube was then transferred to -80°C until mechanical testing began. In total, 171 complete single fibre experiments were performed with 65 from pre-intensified training (slow = 23, fast = 42) and 106 post-intensified training (slow = 33, fast = 73).

On the single fibre testing days, the samples were dissected from bundles in a relaxing solution before being transferred to a 15°C chamber filled with a relaxing solution and tied to pins between a force transducer (model 403A; Aurora Scientific, CAD) and length controller (model 322C; Aurora Scientific) with nylon ties. Average sarcomere length was measured using a high-speed camera and set to 2.8μm. This allows the fibre, upon isometric activation, to shorten to 2.7μm, which is the average optimal sarcomere length for force production in human muscle (39). A fitness contraction was performed in the highest pCa concentration (i.e., pCa=4.5) to ensure that the fibre was not damaged, and ties were secure before testing began. The sarcomere length was remeasured after this step to ensure that it was 2.8μm and readjusted if necessary. The length of the fibre (L_o_) was measured with the microscope, from one suture knot to the other with a reticule. The diameter was measured in 3 places to calculate the cross-sectional area of the individual fibre, assuming circularity.

Prior to recording force, the fibre was placed in a pre-activating solution (reduced Ca^2+^ -buffering capacity) for 10s, then transferred to an activating solution (Ca^2+^ and high [ATP]) for 30s. Seven solutions were used with differing concentrations of Ca^2+^ to assess the force-Ca^2+^ relationship. The concentrations of Ca^2+^ that were used were pCa 7.0, 6.6, 6.4, 6.2, 5.7, 5.5, 4.5, and force measurements were taken at each concentration. A representative tracing can be found in Figure 1B. The maximum force of each fibre was measured as the highest 500-ms moving average along the force-time curve during the time in each pCa solution. After the force was developed, a length step was performed to measure rate of force re-development (k_tr_; Brenner & Eisenberg, 1986). The k_tr_ was measured at each pCa by rapidly shortening the fibre with a ramp of 10 L_o_ /s by 15% of L_o_ and then a rapid (500 L_o_ /s) re-stretch back to L_o_ (Fig. 1B). The rapid shortening step allows for all cross-bridges to break, while the re-stretching step allows for any other cross-bridges to dissociate. The curve of force re-development was fit with a mono-exponential equation y = a (1-e^-kt^) + b to determine k_tr_. Instantaneous stiffness tests were then performed to determine the proportion of attached cross-bridges during a force plateau. This was done by rapidly stretching the fibre (500 L_o_ /s) by 0.3% of L_o_ and dividing the change in force during the stretch by the change in length. Once all tests were performed, there was a final contraction in the activating solution with a pCa concentration of 4.5 to assess if damage to the fibre occurred during contraction. An individual fibre was not included in the final analysis if the force during the final pCa of 4.5 decreased by ≥35% compared to the first experiment with this concentration. Data presented at a pCa of 4.5 was from the first experiment.

### Separation of fibres based on slow and fast phenotype

While MHC isoforms are often used to determine single muscle fibre type for subsequent analysis, we opted to bin fibre type based on fibre phenotype (‘fast’ or ‘slow’) using the k_tr_ values. Previous work has shown that single fibre contractile properties are consistent with MHC isoform, even across muscle groups (18). Furthermore, there are significant differences in mechanical properties between fibre types, justifying that fibre type can be identified based on contractile properties such as shortening velocity (10, 34), or in this case, k_tr_ values. Binning of human fibres has been done previously using unloaded shortening velocity values to separate the fibre types with little error (7% for slow- and 5% for fast-type fibres (10)), providing evidence that mechanical properties can be used as a proxy for biochemical properties (34). Given the scope of the current study, binning based on phenotype is more representative of the mechanical properties of the fibres. Fibres that had a k_tr_ value of less than 40% of the highest k_tr_ from each sample (participant) were labelled as “slow”, and those 40% or higher were labelled “fast”.

### Curve fitting analysis

Each individual joint level force-frequency curve was fit using non-linear regression analysis to obtain the frequency needed to produce 50% (Frequency_50_) of the maximal force pre- and post-intensified training. The Absolute IC50 equation with the least squares regression method was used and the fit was constrained at the top and baseline to 100 and 0, respectively (GraphPad Prism software, version 9.5.1, USA). The curve fitting had an average r^2^ of 0.992±0.001. To estimate the frequency needed to produce 20% (Frequency_20_) and 80% (Frequency_80_) of maximal force, the frequencies were converted into logarithmic values and the “log(agonist) vs. response-Find ECanything” equation was used with the least squares regression method setting the top and bottom constraints to 100 and 0, respectively (GraphPad Prism software, version 9.5.1). The pCa value at which 20%, 50% and 80% of maximal force was produced (i.e., pCa_20_, pCa_50_, and pCa_80_, respectively) were also determined using non-linear regression curve analysis. The “log(agonist) vs. response-Find ECanything equation (pCa_20_, pCa_80_) or log(agonist) vs. normalized response-variable slope (pCa_50_) was used with the least squares regression method setting the top and bottom constraints to 100 and 0, respectively (GraphPad Prism software, version 9.5.1). Single fibres were excluded from the final statistical analysis if either of the following occurred: the force at a pCa concentration of 4.5 decreased by ≥35% at the end of a single experiment and/or the curve fitting produced a r^2^ <0.85. The cut-off criteria of r^2^ <0.85 was decided arbitrarily as we aimed to achieve the largest sample size possible while maintaining quality curve fitting for each fibre analyzed. The total number of fibres excluded for pre-intensified training was 34 (slow = 14, fast = 20) and post-intensified training was 67 fibres (slow = 20, fast = 47) excluded. Thus, the total fibres included in the analysis were as follows: 31 total fibres from 5 participants for pre-intensified training (slow = 9, fast = 22) and 39 total fibres from 10 participants for post-intensified training (slow = 13, fast = 26). The average curve fit was similar pre-intensified training (slow: r^2^=0.962±0.009 and fast: r^2^=0.952±0.006) and post-intensified training (slow: r^2^=0.960±0.009 and fast: r^2^=0.933±0.006).

### Statistical analysis

All statistical analyses were completed in SPSS (IBM, SPSS Statistics, Version 29, USA) and significance was set *a prior* at an α of 0.05. Paired t-tests or repeated measures ANOVA were performed for joint level neuromuscular function testing. Paired t-test effect size was estimated with Cohen’s *d* using the standardized mean difference (*d*_z_). Due to multiple fibres originating from the same participants, linear mixed model analyses were performed for all single fibre data, as has been done previously (5). Participant was set as the random effect and fixed effects were time (pre- and post-intensified training) and muscle fibre phenotype (slow and fast). Where relevant, fixed effect partial eta (η ^2^) values are presented from the linear mixed models. When a significant effect was observed, pairwise comparisons were made with a Bonferroni correction factor. All data is presented as mean ± standard error.

## Results

### Intensified training on joint level neuromuscular function

No differences were observed after 3-weeks of intensified training for joint level MVC force, voluntary activation, twitch force, or twitch half relaxation time (Fig. 2). By contrast, force output was reduced after intensified training during electrically evoked contractions at low frequencies of 5Hz, 8Hz, 10Hz, 15Hz, and 20Hz expressed as both absolute (∼8-27% reduction, all *p≤0.02*) and relative (∼6-26% reduction, all *p≤0.003*) values; (Fig. 3A)). High frequency (i.e., 50 and 100Hz) force output did not change before versus after training (all *p≥0.86*, Fig. 3A). In support of PLFFD at the joint level, there was an increased frequency of stimulation required to evoke the 20% and 50% of the relative force post-intensified training (Frequency_20_: Pre=4.5±0.4Hz versus Post=6.1±0.5Hz, ∼37% difference; Frequency_50_: Pre=11.2±0.6Hz versus Post=13.8±0.8Hz, ∼23% difference), but not the 80% relative force (Frequency_80_: Pre=26.8±1.8Hz versus Post=29.5±1.4Hz, Fig. 3B). Similarly, the ratio of 10:100Hz was reduced after training (Pre=0.47±0.03 versus Post=0.37±0.03, Fig. 3B). These increases resulted in a rightward shift in the force-frequency curve, signifying the presence of PLFFD following a period of intensified training.

**Figure 2.**
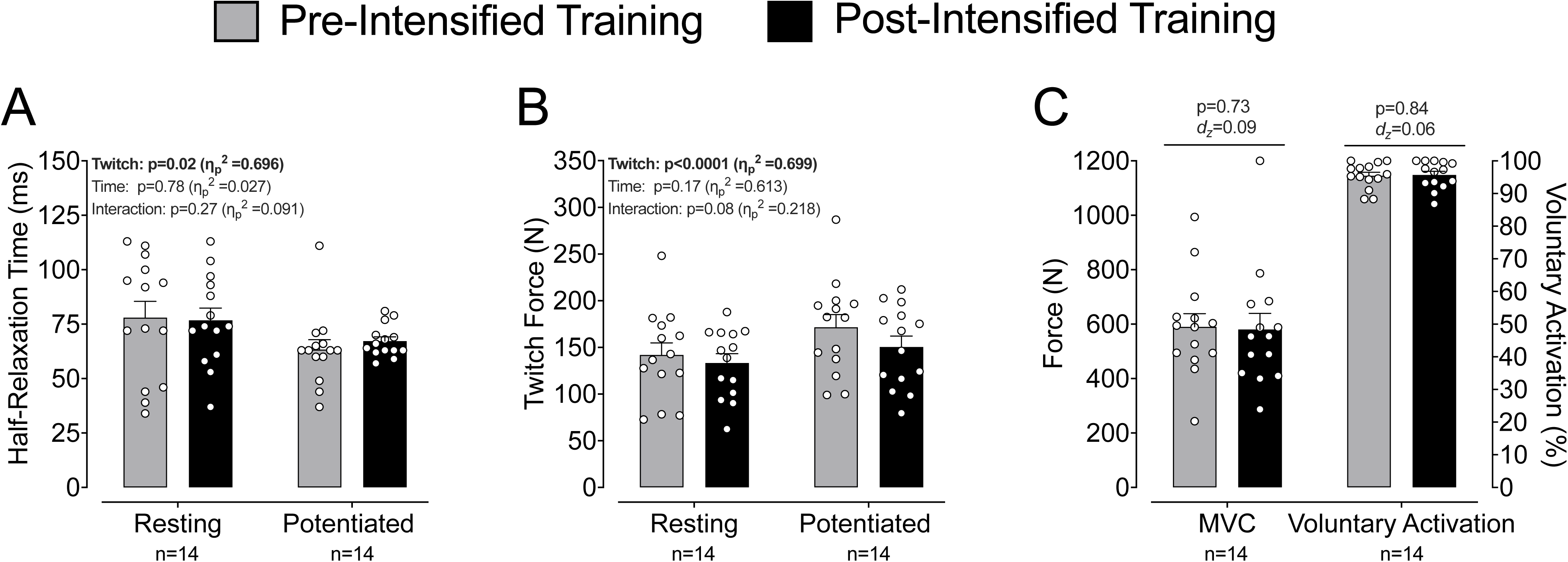
Joint level neuromuscular function tests before (gray bars) and after (black bars) 3-weeks of intensified training. (A) Twitch half-relaxation time. (B) Peak twitch force. (C) Maximal voluntary contraction (MVC) force production and voluntary activation. ANOVA p-values with partial eta (η ^2^) and paired t-test p-values with Cohen’s *d* effect size using the standardized mean difference (*d*_z_) are provided above each outcome. Each circle represents an individual participant. Data is expressed as the mean+standard error of mean (n=14/timepoint).

**Figure 3.**
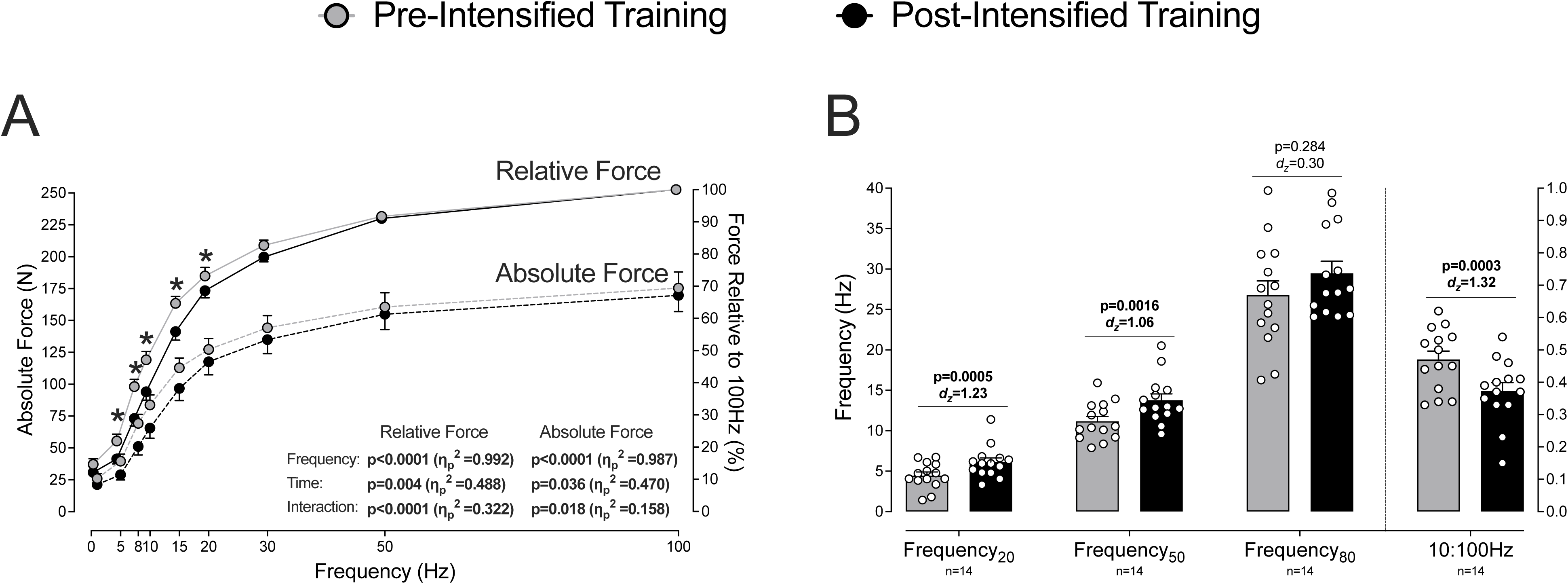
(A) Joint level force-frequency before (gray) and after (black) 3-weeks of intensified training. Data is expressed as an absolute force output (dashed lines) or as a relative percent to the force output to at 100Hz (solid lines). ANOVA p-values with partial eta (η ^2^) for absolute and relative force output are found within the graph. The asterisks represent significant (p<0.05) differences pre-versus post-intensified training for both absolute and relative force outputs at the given frequencies. (B) The frequency required to cause 20% (Frequency_20_), 50% (Frequency_50_), and 80% (Frequency_80_) of the 100Hz force, and the ratio of 10:100Hz force. Paired t-test p-values with Cohen’s *d* effect size using the standardized mean difference (*d*_z_) are provided above each outcome. Data is expressed as the mean+standard error of mean (n=14/timepoint).

### Intensified training on single fibre contractile properties

No significant time × fibre phenotype interactions were observed for peak force nor specific force relative to fibre cross-sectional area; however, fast fibres produced more force than slow fibres (Fig. 4A&B). As expected, based on the binning procedure, fibres categorized as fast had a greater rate of force re-development (k_tr_), than the fibres categorized as slow (Fig. 4C). An effect of time was also observed for the ktr test, such that the single fibres re-developed force faster post-intensified training (Fig. 4C). No differences were observed for instantaneous stiffness across time or phenotype (Fig. 4D).

**Figure 4.**
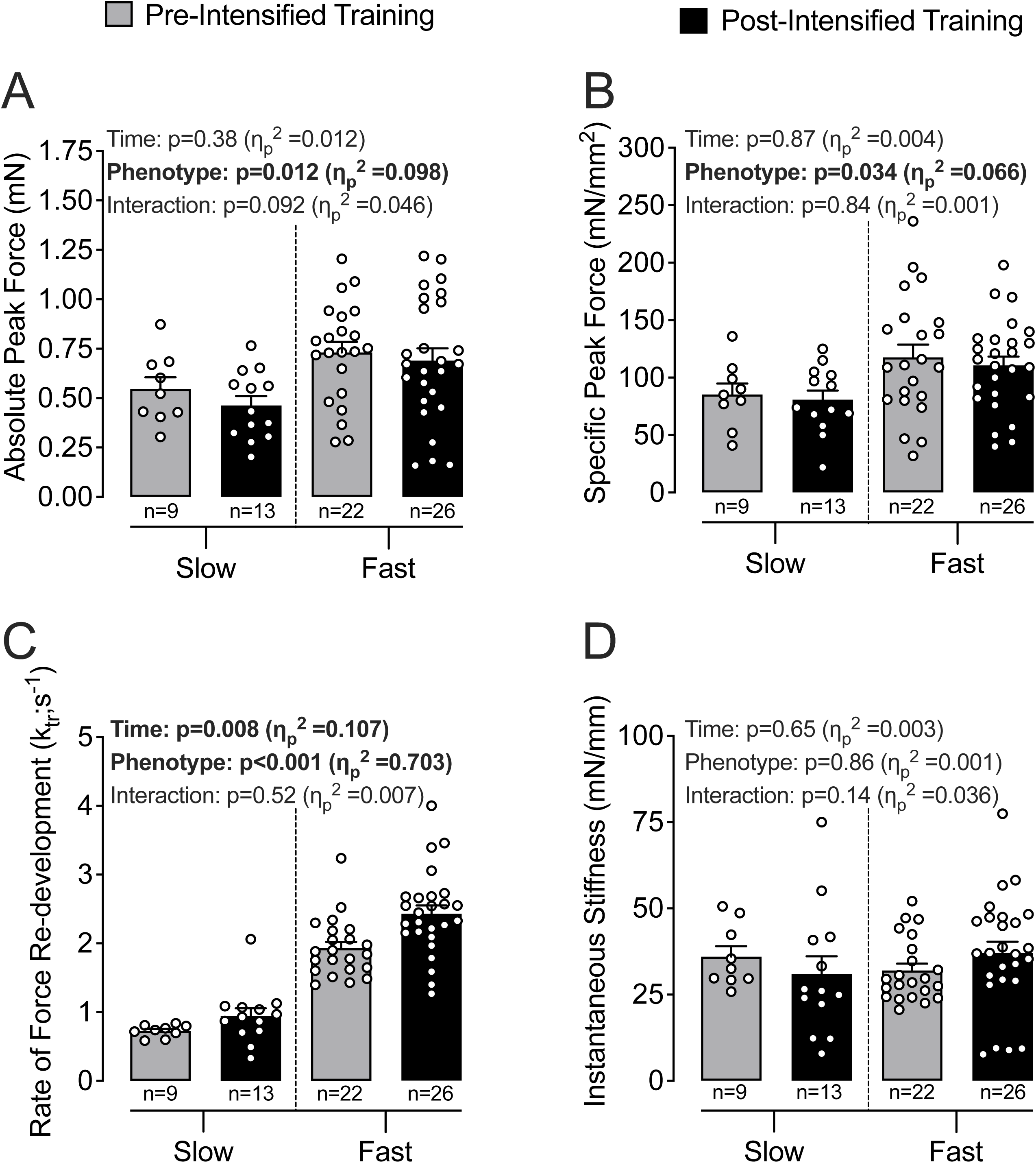
Slow and fast single muscle fibre peak absolute force production (A) or specific force (peak force normalized to cross-sectional area) (gray) and after (black) 3-weeks of intensified training. (C) Rate of force re-development (k_tr_) and (D) instantaneous stiffness at the pCa that evoked the maximal force. Each circle represents an individual single muscle fibre. Linear mixed model effects with partial eta (η ^2^) are provided in each panel. Data is expressed as the mean+standard error of mean (n=9-26/fibre phenotype).

### Intensified training on single fibre myofibrillar Ca^2+^ sensitivity

A main effect of time was observed such that force production was reduced post-intensified training across a range of pCa concentrations regardless of fibre phenotype (Fig. 5A&B). No time × phenotype interaction was observed for markers of myofibrillar Ca^2+^ sensitivity (i.e., pCa_20_, pCa_50_, and pCa_80_ values); however, a main effect of time was observed such that there was a rightward shift in the force-pCa curve with an ∼6% reduction in single fibre pCa_20_, pCa_50_, and pCa_80_ values after intensified training (Fig. 5C-E). Collectively, the data indicates joint level PLFFD after intensified training is driven in part by a reduction in myofibrillar Ca^2+^ sensitivity.

**Figure 5.**
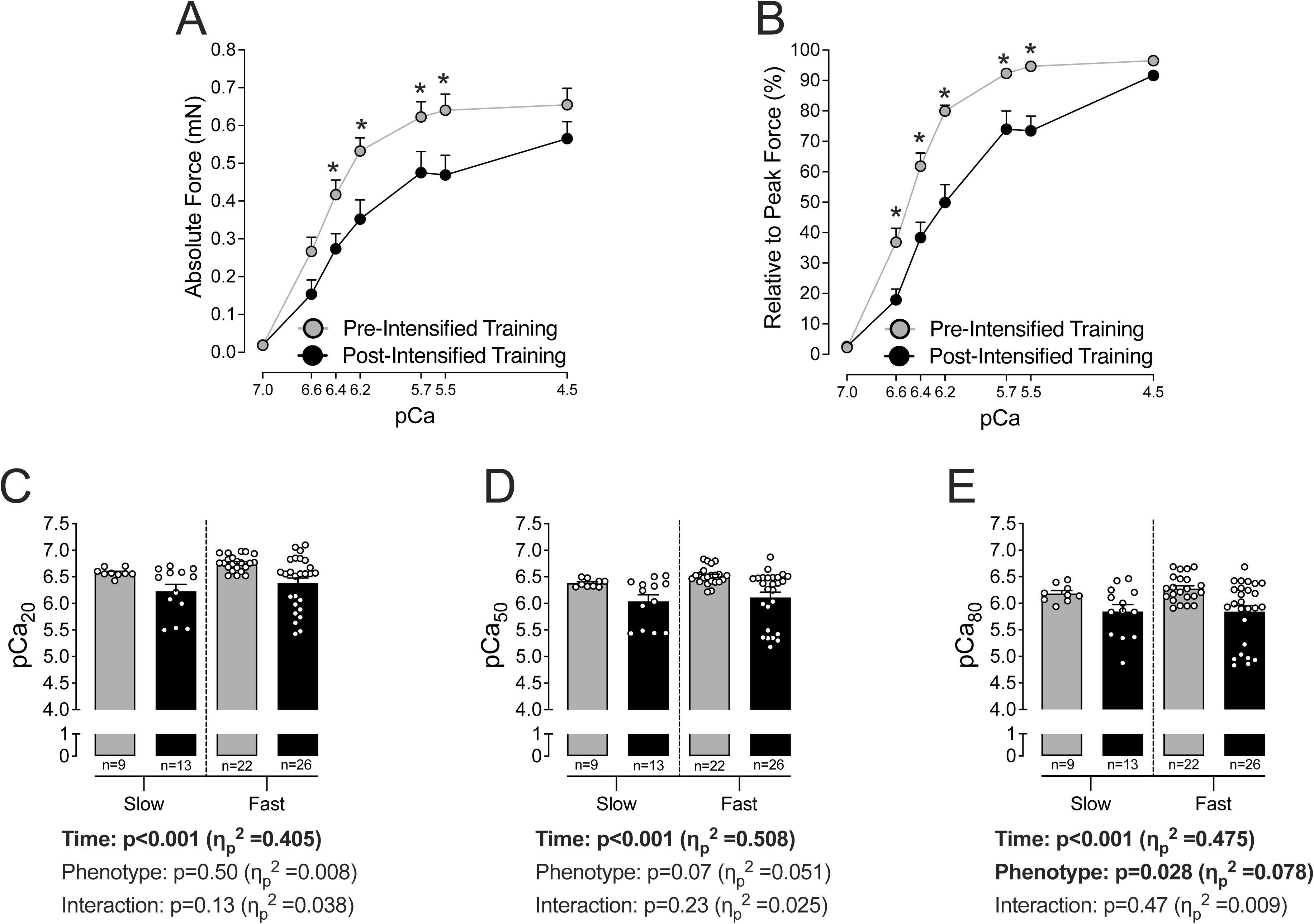
Single muscle fibre level force-calcium curves expressed as absolute force output (A) or as a relative percent to the highest force achieved in each experiment (B). Data from slow and fast fibres are pooled for clarity. The asterisks represent a significant (p<0.05) effect of time when data is pooled across fibre phenotypes in the linear mixed model analysis. The concentration of calcium required to cause (C) 20% of the maximal force output (pCa_20_), (D) 50% of the maximal force output (pCa_50_), and (E) 80% of the maximal force output (pCa_80_) within each force-pCa experiment separated by slow and fast muscle fibre phenotypes. Each circle represents an individual single muscle fibre. Linear mixed model effects and linear mixed model effects with partial eta (η ^2^) for pCa_20_, pCa_50_, and pCa_80_, are provided below each respective panel. Data is expressed as the mean+standard error of mean (n=9-26/fibre phenotype).

## Discussion

The purpose of the present study was to investigate how neuromuscular function is altered following a period of intensified training with insufficient recovery. Pairing joint-level and single muscle fibre-level testing, this is the first study to concurrently characterize the response to intensified training at both macroscopic and microscopic levels. Maximal joint level strength was maintained, but PLFFD was present across frequencies of 5, 8, 10, 15, and 20 Hz, providing evidence that submaximal force production was impaired. At the single muscle fibre level, there was no reduction in maximal force production, but there was a rightward shift in the force-pCa curve driven by a reduction in myofibrillar Ca^2+^ sensitivity, regardless of muscle fibre phenotype. Altogether, impairments in joint level submaximal force production after intensified training may in part be explained by a reduction in myofibrillar Ca^2+^sensitivity in human muscle.

### Maximal force output is not altered at the joint or single fibre level

Maximal strength as assessed via isometric MVC force, has previously been demonstrated to be maintained following an overloading training stimulus (6, 32). In agreement, our data also support that a period of intensified training did not reduce the capacity to voluntarily generate maximal force. In support of this, at the single muscle fibre level, there were also no changes in maximal force. However, we provide novel evidence that intensified training induced PLFFD as relative and absolute force were reduced only at low frequencies (i.e., 5-20 Hz). Notably, joint level PLFFD was observed well after the last exercise bout when exercise-metabolites that impair force production (e.g., inorganic phosphate) were very likely at resting levels. Overall, these results highlight the need to assess force production at submaximal levels as opposed to classical maximal neuromuscular tests such as an isometric MVC to explore mechanisms of whole-body performance decrements during periods of intensified training with insufficient recovery.

### Possible mechanisms driving impairments in joint level submaximal force production

Mechanisms contributing to PLFFD broadly include a reduction in myofibrillar Ca^2+^ sensitivity and reduced sarcoplasmic reticulum Ca^2+^ release (9). The mechanism of PLFFD likely depends on the interaction of the cellular redox state (7, 16, 28, 40), glycogen content (30), and training state (33) of the individual. Given that PLFFD is multifactorial, we intentionally increased exercise duration, intensity, and frequency to determine if myofibrillar Ca^2+^ sensitivity was altered after this period of intensified training. Our findings indicate an ∼6% reduction in pCa_20_, pCa_50_ and pCa_80_ values from pre- to post-intensified training. Thus, a higher concentration of Ca^2+^ (or lower pCa) is required for the same relative force to be produced. Notably, force output at pCa concentrations of 5.7 and 5.5 were reduced from ∼94% of the maximum force pre-trainingto ∼74% of the maximum force after training. Such a reduction in near-maximal force has not been observed for at least a 24h period after high- or low-intensity exercise in elite athletes (17, 21) and after 4-weeks of downhill running in rodents (20). Speculatively, a temporal response may exist where repetitive exercise with inadequate recovery (as done here) decreases force output through multiple mechanisms. From the current data, the reduction in force does not seem to be a result of changes in the number of attached cross-bridges as instantaneous stiffness was unchanged after training. However, it is possible that there may have been a greater proportion of cross-bridges in a weakly bound configuration, analogous to what may occur with aging (24, 34).

### No fibre-type dependency observed

In a *post-hoc* analysis, muscle fibres were binned based on k_tr_ values (e.g., how quickly fibres generate force), and labelled phenotypically as either “slow” or “fast” fibres. This approach bins the fibres based on their actual contractile properties rather than assumed contractile properties based on classic MHC analyses (19). This binning approach has been performed previously, where fibres were binned as slow if their speed was less than 40% of the maximum shortening velocity of the fastest fibre in each group (34). Similar methods have been validated when using contractile properties in place of MHC isoform fibre typing and have been shown to be accurate when compared to MHC isoform classification (10). In this study, we did not find strong evidence to support that intensified training preferentially impaired slow or fast type fibres. This observation contrasts previous work that has shown myofibrillar Ca^2+^ sensitivity is increased in both type I (slow) and II (fast) single muscle fibres after high-intensity exercise (17), and reduced only in type II fibres after 4h of cycling (21). In addition, our single fibre data partly disagrees with a recent study that demonstrated that those with a whole-muscle fast phenotype (measured indirectly based on MRI-derived carnosine content) were more susceptible to impaired running performance after a period of high-volume training (2). The above discrepancies could be due to methodological differences in fibre population identification (e.g., fibre phenotype versus MHC isoform) or phenotype classification (i.e., single muscle fibres versus MRI-derived carnosine content in whole-muscle), but it is also possible that our intensified training program was effective in systematically stressing muscle fibres regardless of their phenotype. Indeed, a difference between the previously mentioned studies and the current study was the purposeful increase in three hallmark exercise variables that facilitate acute stress-responses and training adaptations: training volume, frequency, and intensity that repetitively recruited the entire musculature regardless of fibre type.

Underpinning our observations of joint level PLFFD and impaired single fibre force output is a likely increase in exercise-induced oxidative stress. The depression in submaximal force generation from severely exhaustive exercise can be described by the “inverted U-shape” relationship between the redox state of the muscle and muscle contractile function (9). In support of the oxidative stress hypothesis of PLFFD, myofibrillar Ca^2+^ sensitivity decreases back to baseline when post-exercise single fibres are incubated in the presence of a reducing agent (17) and at least in type II (fast) fibres, may involve redox modifications of a specific cysteine residue in troponin I that is sensitive to changes after acute exercise (28). While increasing cellular oxidative stress can affect contractile function in skeletal muscle, it also has a dual-role of being a key signaling molecule to initiate signaling cascades implicated in training adaptations (as reviewed in (35)). Therefore, a necessary consequence of intensified training with the goal of an overload stimulus is a decrement in muscle contractile function but, with adequate recovery, the training adaptations acquired during this period can be realized to increase sport performance.

### Study Considerations

Since the change in the single fibre data for pCa_20_, pCa_50_, and pCa_80_ values were smaller (∼6%) than joint-level PLFFD as inferred through changes in the Frequency_20_ (23%) and Frequency_50_ (37%), our findings do not exclude other mechanisms besides changes in myofibrillar Ca^2+^ sensitivity such as impaired sarcoplasmic reticulum Ca^2+^ release contributing to impaired joint level performance. In addition, the use of non-physiological conditions (e.g., temperature or metabolite concentrations) that do not reflect those present during intense exercise could limit the extrapolation of these findings to sport performance. Lastly, the lack of a control group performing their regular training throughout the duration of the study prevents certainty that the observed findings were a direct result of the intensified training program.

## Conclusion

Intensified training with insufficient recovery can reduce sport performance, and it can take days to weeks for these functional impairments to be ameliorated (27). The findings from the present investigation demonstrate a novel mechanism of performance reduction following overload training and has important implications for athletic training monitoring and recovery strategies. Our data indicate intensified training induces PLFFD due in part to reductions in myofibrillar Ca^2+^ sensitivity.

## Abbreviations

L_o_: length of the fibre
MVC: maximum voluntary contraction
PLFFD: prolonged low frequency force depression
[Ca^2+^_i_]: free Ca^2+^ concentration
SOD2: superoxide dismutase 2

## Author contributions

Conception and study design: CP, AMC, GAP, AJC, JFB; Participant recruitment and scheduling: AMC; Data collection and/or analysis: OPR, CP, EFH, GAP, AMC; Initial manuscript draft and interpretation: OPR, CP, GAP; Manuscript interpretation and revision: OPR, CP, EFH, AMC, AJC, JFB, GAP; All authors have approved the final version of the manuscript.

## Acknowledgements

We would like to thank all the participants in this study. Figure 1 was in part created with BioRender.com and are published with permission.

## Supplemental material available at

DOI: https://doi.org/10.6084/m9.figshare.24231727.v2

URL: https://figshare.com/articles/journal_contribution/Supplementary_information/24231727

## Data accessibility

Supporting data are available upon request.

